# Rapid neural categorization of facelike objects predicts the perceptual awareness of a face (*face pareidolia*)

**DOI:** 10.1101/2021.02.19.431826

**Authors:** Diane Rekow, Jean-Yves Baudouin, Renaud Brochard, Bruno Rossion, Arnaud Leleu

## Abstract

The human brain rapidly and automatically categorizes faces *vs*. other visual objects. However, whether face-selective neural activity predicts the subjective experience of a face – *perceptual awareness* – is debated. To clarify this issue, here we use *face pareidolia*, i.e., the illusory perception of a face, as a proxy to relate the neural categorization of a variety of facelike objects to conscious face perception. In Experiment 1, scalp electroencephalogram (EEG) is recorded while pictures of human faces or facelike objects – in different stimulation sequences – are interleaved every second (i.e., at 1 Hz) in a rapid 6-Hz train of natural images of nonface objects. Participants do not perform any explicit face categorization task during stimulation, and report whether they perceived illusory faces post-stimulation. A robust categorization response to facelike objects is identified at 1 Hz and harmonics in the EEG frequency spectrum with a facelike occipito-temporal topography. Across all individuals, the facelike categorization response is of about 20% of the response to human faces, but more strongly right-lateralized. Critically, its amplitude is much larger in participants who report having perceived illusory faces. In Experiment 2, facelike or matched nonface objects from the same categories appear at 1 Hz in sequences of nonface objects presented at variable stimulation rates (60 Hz to 12 Hz) and participants explicitly report after each sequence whether they perceived illusory faces. The facelike categorization response already emerges at the shortest stimulus duration (i.e., 17 ms at 60 Hz) and predicts the behavioral report of conscious perception. Strikingly, neural facelike-selectivity emerges exclusively when participants report illusory faces. Collectively, these experiments characterize a neural signature of face pareidolia in the context of rapid categorization, supporting the view that face-selective brain activity reliably predicts the subjective experience of a face from a single glance at a variety of stimuli.

**Highlights:**

- EEG frequency-tagging measures the rapid categorization of facelike objects
- Facelike objects elicit a facelike neural categorization response
- Neural face categorization predicts conscious face perception across variable inputs

## 1. Introduction

Humans are very good and fast at categorizing visual stimuli as faces (e.g., Crouzet et al., 2010; Hershler et al., 2010; Scheirer et al., 2014). At the neural level, this critical face categorization function is subtended by a large network of cortical areas in the ventral occipito-temporal cortex (VOTC) (e.g., Gao et al., 2018; Jonas et al., 2016; Sergent et al., 1992; Zhen et al., 2015; Grill-Spector et al., 2017 for review) and leads to specific signatures in scalp electroencephalography (EEG) (e.g., Bentin et al., 1996; Jeffreys, 1996; Rossion et al., 2015). However, whether face-selective neural activity predicts the subjective experience of a face – *perceptual awareness*, is debated (Aru et al., 2012; Harris et al., 2011; Moutoussis & Zeki, 2002; Navajas et al., 2013; Perry, 2016; Philiastides & Sajda, 2006; Retter et al., 2020; Tanskanen et al., 2007; Tong et al., 1998). This is essentially due to the challenge of measuring face categorization in the brain, i.e., measuring a neural response that incorporates high selectivity to faces (*vs*. many nonface categories) and generalizability across a wide range of variable face stimuli (e.g., Rossion, 2014 for a discussion).

Recently, a valid measure of rapid and automatic face categorization has been developed in scalp EEG (e.g., Jacques et al., 2016; Retter & Rossion, 2016; Rossion et al., 2015), neuroimaging (Gao et al., 2018), and intracranial recordings (Hagen et al., 2020; Jonas et al., 2016) using a frequency-tagging approach (Norcia et al., 2015 for review). With this approach, a rapid stream of forward- and backward-masked natural images of many living and non-living objects is presented at a base rate (e.g., 6 Hz) and variable exemplars of human faces are interspersed at a lower rate (e.g., 1 Hz) while participants do not explicitly categorize faces (Fig. 1A & 1B). This paradigm thus dissociates two brain responses automatically elicited at predefined frequencies: a general visual response (base rate) and a face categorization response (face presentation rate). The general response reflects the neural activity elicited by the fast train of stimuli, while the face categorization response captures face-selective activity generalized across face stimuli. This latter response is large, sensitive, reliable, and not accounted for by the amplitude spectrum of the images (Gao et al., 2018; Rossion et al., 2015). Albeit sporadic (i.e., recorded for only a subset of faces), the face categorization response is already fully elicited (i.e., of full magnitude) at very high speed of stimulation (i.e., at 60 Hz; stimulus duration: 17 ms) and saturates (i.e., becomes systematic) at 12 Hz (stimulus duration: 83 ms; Retter et al., 2020). Importantly for our purpose, these variable occurrences of the response as a function of stimulus duration are associated with behavioral reports of face perception, revealing that rapid and all-or-none face categorization in the brain predicts perceptual awareness (Retter et al., 2020).

**Figure 1.**
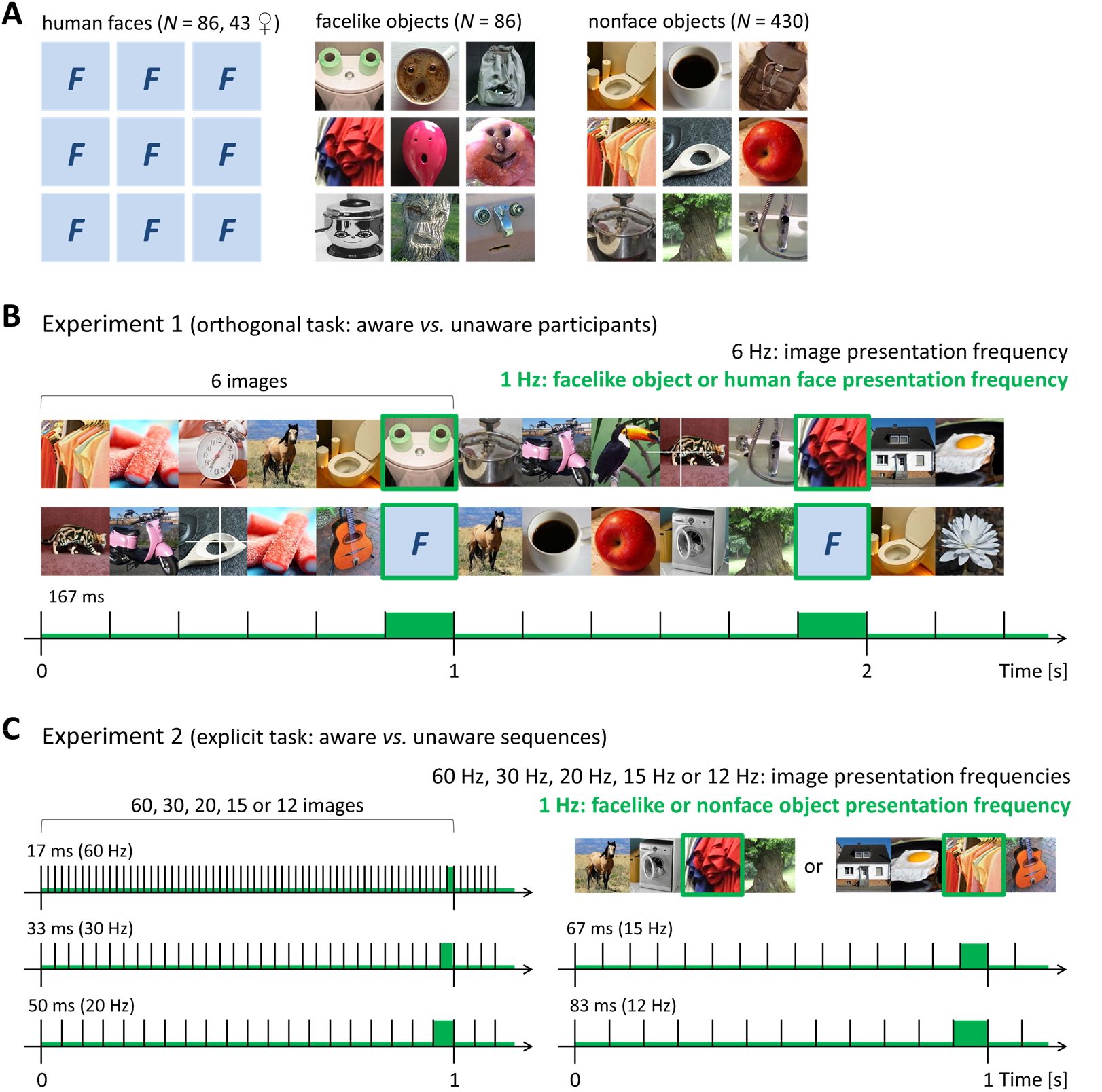
Measuring face pareidolia with EEG frequency-tagging. **A**. Examples of variable natural images of human faces (indicated here by placeholders *F*), facelike objects and nonface objects used as stimuli in both experiments. **B**. Example of ≈ 2 seconds of visual stimulation at a 6-Hz presentation frequency (i.e., 6 images per second without inter-stimulus interval, stimulus duration ≈ 167 ms) used in Experiment 1. Facelike objects or human faces are inserted at 1 Hz (i.e., every 6^th^ stimulus) in dedicated sequences. This procedure tags two brain responses in the EEG frequency spectrum: a general response (6 Hz and harmonics, i.e. integer multiples) to all visual cues rapidly changing at 6 Hz, and a categorization response (1 Hz and harmonics) reflecting the visual categorization of facelike objects or human faces (i.e., discrimination from nonface objects and generalization across exemplars). Participants perform a cross-detection task and are asked post-stimulation if they noticed illusory faces to compare perceptually aware and unaware participants. **C**. Examples of ≈ 1 second of stimulation at 5 presentation frequencies (60 Hz, 30 Hz, 20 Hz, 15 Hz, 12 Hz; stimulus durations: 17 ms, 33 ms, 50 ms, 67 ms, 83 ms) used in Experiment 2. Facelike or nonface objects are interspersed at 1 Hz in dedicated sequences. This time, participants explicitly attend to the stimuli and report after each sequence if they perceived illusory faces to compare sequences associated with awareness or not.

Despite the large variability of the face stimuli used in frequency-tagging studies, they all show clear human facial features, are all recognized as faces by human observers, and very likely to be recognized as faces with high accuracy by an artificial system (e.g., Scheirer et al., 2014; see Grill-Spector et al., 2018 for a discussion). However, an in-depth understanding of conscious face perception from rapid neural categorization must consider that face percepts also emerge from a variety of inputs in the natural visual environment despite the absence of human facial features, namely *face pareidolia* (see examples in Fig. 1A). Prior studies have documented how face pareidolia elicit activity within face-selective regions in the VOTC (Dolan et al., 1997; Hadjikhani et al., 2009; Kanwisher et al., 1998; McKeeff & Tong, 2007; Rossion et al., 2011), or a facelike EEG response over right occipito-temporal scalp sites (Caharel et al., 2013; Churches et al., 2009; Sagiv & Bentin, 2001). Facelike neural activity is generally identified when stimuli are consciously reported as faces by human observers (Andrews & Schluppeck, 2004; Bentin et al., 2002; George et al., 2005; Shafto & Pitts, 2015), contrary to stimuli judged as facelike by a computational face-detection system (Moulson et al., 2011). Nevertheless, previous studies have not clarified whether a large set of naturalistic facelike objects can be rapidly and automatically categorized in a train of masked stimuli depicting the same object categories, how similar is neural facelike categorization to the categorization of genuine human faces, and whether such a rich categorization response predicts the conscious report of illusory faces in individual participants.

To fill this gap in knowledge, here we employ EEG frequency-tagging in two experiments to provide a neural categorization response reflecting the conscious perception of a face in a large set of naturalistic facelike stimuli contrasted to nonface stimuli depicting similar objects (Fig. 1A). The nonface stimuli are presented at the base rate and the facelike stimuli at a lower rate, such that a facelike categorization response emerges only if stimuli depicting similar objects (i.e., facelike and nonface stimuli) elicit dissimilar neural activity, whereas stimuli depicting dissimilar objects (i.e., facelike stimuli) elicit similar neural activity according to their facelikeness (Fig. 1B & 1C). In *Experiment 1*, we present 27 participants with 40-second-long sequences at a 6-Hz base rate (i.e., 6 images per second, stimulus duration ≈ 167 ms) and facelike objects or human faces are interleaved every 6 stimuli (i.e., at 1 Hz; Fig. 1B) to tag the categorization of illusory and human faces at 1 Hz and harmonics (i.e., integer multiples) and estimate the facelikeness of the former. Importantly, participants perform an orthogonal cross-detection task ensuring implicit exposure to facelike stimuli. They are then queried post-stimulation whether they noticed facelike objects, and classified as perceptually aware or unaware participants. In *Experiment 2*, another 22 participants are presented with 16-second-long sequences at 5 different base rates (60 Hz, 30 Hz, 20 Hz, 15 Hz, 12 Hz), such that stimulus duration varies from 17 to 83 ms as in Retter et al. (2020). Facelike or nonface objects are always interspersed at 1 Hz in dedicated sequences (i.e., facelike objects in half of the sequences). Contrary to Experiment 1, participants are informed of the presence of facelike objects before testing and must report after each sequence if they perceived illusory faces to contrast sequences associated with perceptual awareness or not in each participant. Overall, through these two experiments, we demonstrate that a facelike categorization response to a wide range of facelike stimuli already emerges at a short 17-ms stimulus duration in the human brain, and predicts conscious face perception in individual participants.

## 2. Materials and Methods

### 2.1. Experiment 1

#### 2.1.1. Participants

Twenty-seven participants (12 females, 6 left-handed (3 females), mean age: 22.5 ± 2.9 (*SD*) years, range: 19–31 years) took part in the experiment and were compensated for their participation. All reported normal/corrected-to-normal visual acuity, and none reported a history of neurological/psychiatric disorder. They provided written informed consent prior to the experiment. Testing was conducted in accordance with the Declaration of Helsinki and approved by the French ethics committee (CPP Sud-Est III - 2016-A02056-45).

Since participants performed an orthogonal task, we asked them three questions post-stimulation to determine whether they perceived facelike objects. First, we asked if they noticed something particular during the experiment. If they did not mention illusory faces, we then asked whether they noticed something about the stimuli. Note that all participants reported here the presence of human faces but none detected their periodicity. Again, if participants did not mention illusory faces, we finally questioned them about the presentation of facelike objects. Based on this interview, participants were split in two groups, one group that mentioned illusory faces in at least one question (i.e., perceptually *aware* participants, *N* = 13, 5 females, 2 left-handed (1 female), mean age: 23.2 ± 3.5 years, range: 19– 31 years), and another group that did not (i.e., *unaware* participants, *N* = 14, 7 females, 4 left-handed (2 females), mean age: 21.9 ± 2.3 years, range: 19–27 years). The two groups did not significantly differ in age (*T*25 = 1.21, *p* = .24), sex (*X*^2^1 = .55, *p* = .36), and handedness (*X*^2^1 = .41, *p* = .68).

#### 2.1.2. Stimuli

Stimuli were color natural images of 86 human faces (43 females), 86 facelike objects and 430 nonface objects cropped to a square and sized to 300 × 300 pixels. All stimuli were embedded in their original scenes and differed in size, viewpoint, lighting and background so that their physical characteristics were widely variable (examples in Fig. 1A, full set available upon request from the authors). In addition, human faces varied largely in age, sex, race and expression. Face and nonface images were adapted from previous studies (e.g., Jacques et al., 2016; Retter & Rossion, 2016; Rossion et al., 2015) or collected from the Internet. Nonface objects were various biological and manufactured objects with several exemplars (i.e., between 3 and 20) in each category (listed in *Supplementary Materials and Methods*). Facelike images were selected among a large set of 224 pictures collected from the Internet when searching for ‘face pareidolia’. Selection was made according to the images judged as the most facelike in a pretest (*Supplementary Materials and Methods* & Fig. S1). Critically, facelike images depicted various object categories (between 1 and 5 exemplars in each category) matching some of those used for nonface objects (listed in *Supplementary Materials and Methods*). Hence, facelike objects differed from nonface objects only in their overall facelike appearance (Fig. 1A).

Face and facelike stimuli were both divided in two sets of 43 pictures. For human faces, one set contained 22 females and the other one 21 females. For facelike images, at least one exemplar of each object category was allocated to each set. These two sets ensured that all face and facelike stimuli were presented to every participant. During the experiment, stimuli were displayed at the center of a 24-inch LED screen (60 Hz refresh rate, resolution: 1920 × 1080 pixels) on a mid-level grey background (i.e., 128/255 in greyscale). From a viewing distance of 57 cm, they subtended approximately 8.3° of visual angle.

#### 2.1.3. Procedure

The procedure was adapted from previous face categorization experiments using EEG frequency-tagging (e.g., Jacques et al., 2016; Retter & Rossion, 2016; Rossion et al., 2015). Images were presented at a fast base rate of 6 Hz (i.e., 6 images per second, ≈ 167 ms per image) without inter-stimulus interval (forward- and backward-masking; Fig. 1B). In each stimulation sequence, nonface objects were used as base stimuli. In different sequences, human faces or facelike objects were periodically inserted every 6^th^ stimulus (i.e., at 6/6 = 1 Hz; 1 second between two human faces or facelike objects).

After electrode-cap placement, participants were seated in a light- and sound-isolated cabin in front of the stimulation screen. Their head was maintained on a chinrest at a distance of 57 cm from the screen. Stimulation sequences started with a 2-second fade-in of increasing contrast modulation depth (0 to 100%), followed by the full-contrast stimulation lasting 40 seconds and then followed by a 2-second fade-out of decreasing contrast modulation depth (100 to 0%). Both fade-in and fade-out were used to reduce eye-blinks and movements elicited by the sudden onset or offset of flickering stimuli. Sequences were flanked by variable pre- and post-stimulation intervals of 0.5–1.5 seconds of uniform grey background. For both face and facelike stimuli, each set of 43 images was used in half of the stimulation sequences while the 430 nonface objects were used in all sequences. Each experimental condition (i.e., category at 1 Hz) was repeated 6 times (i.e., 3 times for each stimulus set), resulting in 12 sequences throughout the experiment. They were divided in 3 blocks of 4 sequences, each block presenting two sequences per condition (i.e., one per stimulus set). Blocks and sequences within blocks were randomly presented across participants. In each sequence, stimuli were randomly selected from their respective sets.

#### 2.1.4. Orthogonal behavioral task

An orthogonal behavioral task was designed to ensure that participants were exposed to the stimulation. During each sequence, they were asked to detect 8 brief (200 ms) random appearances of a 300 × 300 pixels large white cross on the images by pressing the spacebar of a keyboard with both index fingers as quickly as possible. A minimum interval of 2 seconds was introduced between two cross-onsets. Both accuracy and RTs for correct detections (ranging between 100 and 1000 ms) were submitted to a repeated-measures ANOVA with *Category* (human faces *vs*. facelike objects) as a within-subject factor and *Group* (aware *vs*. unaware) as a between-subject factor.

#### 2.1.5. EEG acquisition and preprocessing

Scalp electroencephalogram (EEG) was continuously acquired from a 64-channel BioSemi Active-Two amplifier system (BioSemi, The Netherlands) with Ag/AgCl electrodes located according to the 10–10 classification system. During recording, the Common Mode Sense (CMS) active electrode was used as reference and the Driven Right Leg (DRL) passive electrode was used as ground. Electrode offset was held below ± 15 μV for each electrode and EEG was sampled at 1024 Hz.

All EEG analyses were carried out in Letswave 6 (https://www.letswave.org/) running on Matlab 2017 (MathWorks, USA). For each participant, continuous datasets were first bandpass filtered at 0.1– 100 Hz using a Butterworth filter (4^th^ order) and then downsampled to 256 Hz. Datasets were segmented into 45-second epochs for each stimulation sequence (12 per participant, 2 conditions × 6 repetitions), including 1 second before the fade-in and 1 second after the fade-out. An Independent Component Analysis (ICA) with a square mixing matrix was computed (Makeig et al., 1996) to isolate and remove components corresponding to eye-blinks (i.e., one component recorded over Fp channels per participant) and to additional artifacts recorded over frontal and temporal channels (mean number across participants: 2.3, range: 0–4, no significant difference between groups of participants, *T25* = 0.91, *p* = .37). Remaining noisy or artifact-ridden channels were replaced using linear interpolation from the 4 neighboring channels (mean number across participants: 0.9, range: 0–5, no significant difference between groups of participants, *T*25 = 0.16, *p* = .88). EEG epochs were then re-referenced to the average of the 64 channels.

#### 2.1.6. EEG frequency-domain analysis

In line with previous face categorization studies (e.g., Jacques et al., 2016; Retter & Rossion, 2016; Rossion et al., 2015), our paradigm was designed to tag two different brain responses at two predefined frequencies within a single stimulation sequence, and to quantify them in the EEG amplitude spectrum using frequency-domain analysis: (1) a general response at 6 Hz and harmonics (i.e., integer multiples) elicited by the stream of images (i.e., both nonface and face/facelike images) and capturing the visual response to all cues (e.g., local contrast) rapidly changing 6 times per second; (2) a categorization response at 1 Hz and harmonics reflecting the *differential* response to face or facelike stimuli. Thanks to the rapid and periodic mode of stimulation, this response indexes single-glance visual categorization of human faces and facelike objects implying discrimination from nonface objects and generalization across category exemplars despite widely variable images. It is not accounted for by the amplitude spectrum of the stimuli (Gao et al., 2018; Rossion et al., 2015) and is immune to temporal predictability elicited by periodicity (Quek & Rossion, 2017). Note that the 1-Hz rate of face or facelike presentation allows enough time between image-onsets (i.e., 1 second) for the full face categorization response to develop (≈ 450 ms in duration, Retter & Rossion, 2016).

For each participant, the 6 epochs recorded for each condition were averaged to reduce EEG activity non-phase-locked to the stimuli, thus resulting in a single 45-second epoch per condition. Epochs were then precisely cropped from the onset of the full-contrast stimulation to 40 seconds so as to contain an exact integer number of 1-Hz cycles (i.e., 40 cycles, 10240 time bins). A fast Fourier transform (FFT) was applied to every epoch and amplitude spectra were extracted for all channels with a high frequency resolution of 1/40 = 0.025 Hz. Thanks to this high resolution, 40 frequency bins were extracted every 1-Hz step, allowing unambiguous identification of the tagged brain responses and estimation of noise amplitude from surrounding frequency bins. Given our objective to identify a selective response to facelike objects that reflects their categorization as faces, we considered the EEG data recorded for sequences containing human faces as a reference to determine the range of harmonics (i.e., tagged frequencies and their integer multiples) and regions-of-interest (ROIs) for further analysis.

The range of harmonics (Table S1) for the brain responses to the 6-Hz stimulation and the 1-Hz face presentation was defined from the FFT amplitude spectra averaged across channels and participants (Fig. 2A). *Z*-scores were calculated as the difference between the amplitude at the target frequency bin and the mean amplitude of the surrounding noise (≈ ± 0.3 Hz: estimated from 20 frequency bins, 10 on each side, excluding the immediately adjacent and the 2 most extreme (minimum and maximum) bins) divided by the standard deviation of the noise. Harmonics were included until *Z*-scores were no longer significant (i.e., *Z* > 1.64, *p* < .05, one-tailed, signal > noise). For the general response, all harmonics were significant (i.e., 8 harmonics, from 6 Hz to 48 Hz, harmonics were not considered beyond the 50 Hz response elicited by AC power). For the face categorization response, harmonics reached significance up to 26 Hz (i.e., 26^th^ harmonic). The overall responses were then condensed by summing amplitudes across significant harmonics (excluding harmonics corresponding to the general response (i.e., 6 Hz, 12 Hz, 18 Hz, 24 Hz) for the categorization response) for each category, channel and participant. Henceforth, mentions of the general and categorization responses to either human or illusory faces will refer to these amplitudes summed across harmonics.

**Figure 2.**
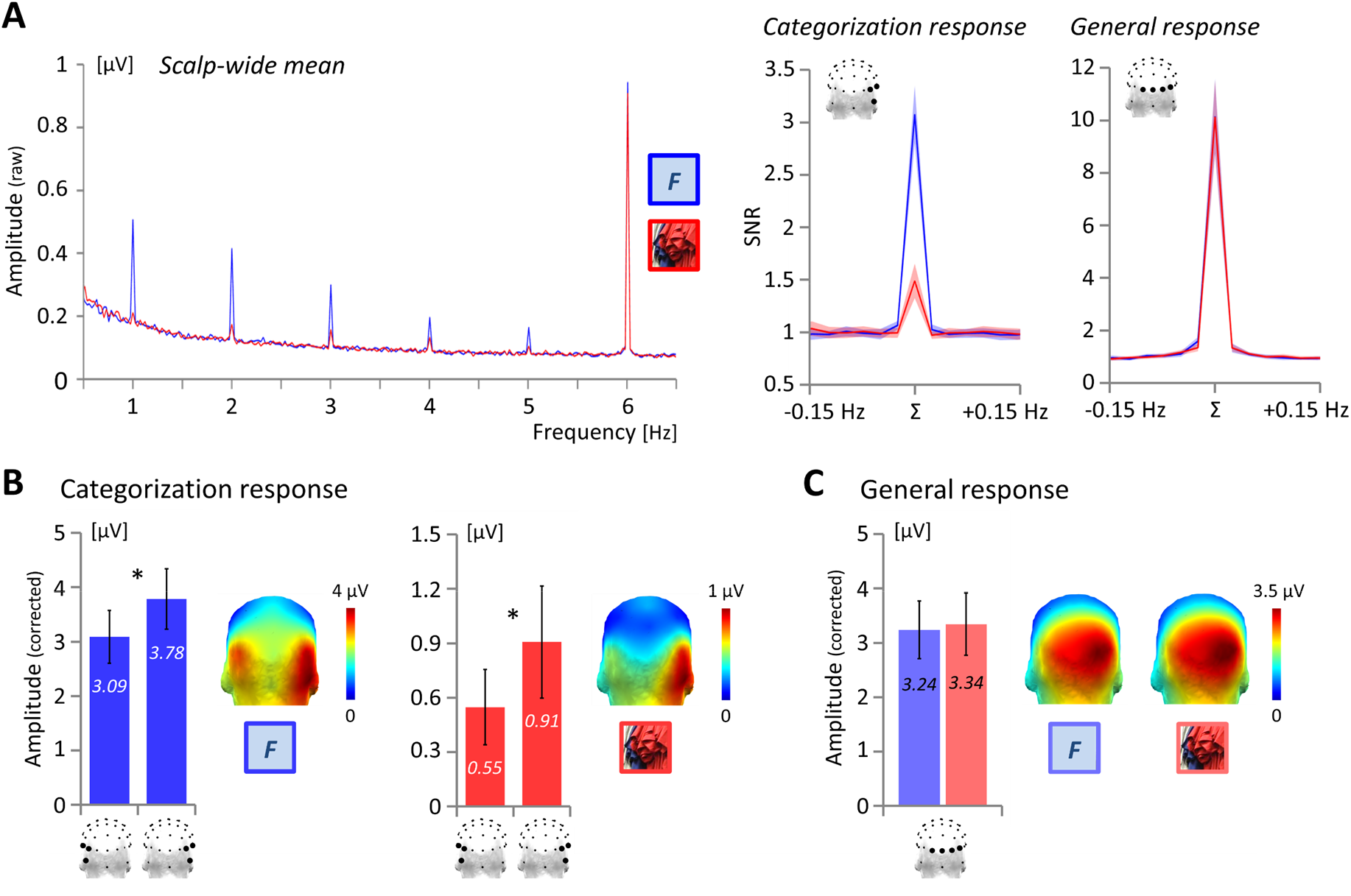
Brain responses elicited in sequences presenting human faces (blue) or facelike objects (red) among nonface objects. **A**. Left: Grand-averaged FFT raw amplitude spectra (across 64 channels). Both types of stimulation sequences elicit a large response at the 6-Hz image presentation frequency (i.e., general response to all images). Though stronger to human faces, responses are clearly visible (i.e., of higher amplitude than surrounding frequency bins) at the 1-Hz human/illusory face presentation frequency (i.e., categorization response) and its harmonics (i.e., integer multiples, here 2 Hz, 3 Hz, 4 Hz, 5 Hz). Right: Grand-averaged signal-to-noise ratio (SNR) of the categorization and general responses over the right occipito-temporal and middle occipital regions, respectively. Responses are summed across significant harmonics (Σ) and compared to surrounding frequencies (± 0.15 Hz, SNR ≈ 1, signal ≈ noise). SNR is high for both responses (categorization response: SNR ≈ 3 and 1.5 for human faces and facelike objects, respectively; general response: SNR ≈ 10 for both categories). **B**. Grand-averaged summed corrected amplitude of the categorization responses over the left and right occipito-temporal regions (* *p* < .05). **C**. Grand-averaged summed corrected amplitude of the general response over the middle occipital region. For both categorization (B) and general (C) responses, topographies are illustrated by head maps (posterior view). Shaded areas in (A) and error bars in (B) and (C) represent 95% confidence intervals.

To quantify the magnitude of each brain response in a value expressed in microvolts (μV), we isolated the response from noise level by subtracting out the mean amplitude of the surrounding frequency bins, leading to notional corrected amplitudes of zero in the absence of response. Corrected amplitudes were used to define the ROIs, and the ROIs were used to conduct group-level statistical analyses and significance estimation of individual brain responses. The strength of each brain response within ROIs (Fig. 2B & 2C) was also estimated with signal-to-noise ratios (SNR) computed by dividing the uncorrected response amplitude by the mean surrounding noise.

In line with previous studies (Jacques et al., 2016; Rossion et al., 2015), both categorization and general responses in the human face condition present a right hemisphere advantage, but the categorization response is laterally distributed over occipito-temporal regions (Fig. 2B) while the general response is located over the middle occipital cortex (Fig. 2C). Accordingly, we defined two symmetrical occipitotemporal ROIs for the face categorization response and one middle occipital ROI for the general visual response by considering the channels with the maximal group-level corrected amplitudes. For the face categorization response, the largest amplitude was observed over channel P10 (4.27 μV), followed by PO8 (3.81 μV), PO7 (3.37 μV), P8 (3.27 μV), P9 (2.96 μV) and P7 (2.93 μV). We thus defined homologous right and left occipito-temporal ROIs (respectively rOT and lOT), each comprising 3 contiguous channels (r/lOT: P10/9, PO8/7, P8/7). For the general visual response, the strongest amplitudes were recorded over channels O2 (3.50 μV), PO8 (3.32 μV), Oz (3.10 μV) and O1 (3.04 μV). The single middle occipital ROI thus encompassed these 4 neighboring channels.

Statistical analyses were conducted separately for the categorization and general responses. In addition, for each response, two separate analyses were also consecutively performed. The first one evaluated the response elicited by facelike objects compared to human faces with the whole participant sample for an initial characterization of the neural signature of illusory face perception. We ran a repeated-measures ANOVA on individual corrected amplitudes with *Category* (human faces *vs*. facelike objects) as a within-subject factor. The within-subject factor *Hemisphere* (rOT *vs*. lOT) was also included for the categorization response. The second analysis aimed at determining whether the neural patterns identified in the first analysis depend on the perceptual awareness of illusory faces. For this purpose, repeated-measures ANOVAs were conducted for each stimulus category with *Group* (aware *vs*. unaware) as a between-subject factor. In all analyses, post-hoc comparisons were conducted for significant effects using *T*-tests and the false discovery rate (FDR) procedure was applied to adjust *p-* values for multiple comparisons (Benjamini & Hochberg, 1995). Since corrected amplitudes should not differ from zero in the absence of response, significance of the grand-averaged brain responses was estimated by identifying whether the 95% confidence interval (*CI*, calculated across participants) around the mean response amplitude did not include zero.

Finally, two other analyses were carried on to estimate the significance of the brain responses in every individual participant and to determine whether the topographies of the categorization responses to facelike objects and human faces are reliable and comparable. For the first analysis, the significance of individual responses within ROIs was estimated using *Z*-scores (see above). For the second analysis, the 6 epochs (i.e., time series) recorded for sequences presenting human faces or facelike objects were split according to stimulus sets, resulting in 2 × 3 epochs for each condition. Epochs were then averaged and following processing steps were similar to those previously described in order to isolate both general and categorization responses to either human faces or facelike objects expressed in summed corrected amplitudes separately for each stimulus set. After grand-averaging individual responses, we computed Pearson’s correlations between the categorization responses obtained for each set using the 64 channels as observations. We thus estimated the topographical reliability of the categorization response across stimulus sets for both categories. Correlations were also calculated between both categorization responses to determine whether their scalp distributions are close. As a control index expected to reveal a lower topographical similarity, the correlation between the categorization response and the general response recorded for sequences containing facelike objects was finally computed.

### 2.2. Experiment 2

#### 2.2.1. Participants

We tested 22 participants (15 females, 1 left-handed (female), mean age: 21.4 ± 4 (*SD*) years, range: 18–33 years) who did not participate in Experiment 1. They reported normal/corrected-to-normal visual acuity and no history of neurological/psychiatric disorder. They provided written informed consent prior to the experiment and were compensated. Testing was conducted in accordance with the Declaration of Helsinki and approved by the French ethics committee (CPP Sud-Est III - 2016-A02056-45).

#### 2.2.2. Stimuli

Stimuli were the 80 facelike objects judged as the most facelike in the pretest conducted before Experiment 1 (*Supplementary Materials and Methods* & Fig. S1) and the 430 nonface objects used in Experiment 1 (examples in Fig. 1A). An additional set of 15 facelike images (judged as the most facelike after the 80 first ones) was also used before testing to illustrate which kind of stimuli participants must detect (see *Explicit behavioral task*). The 80 facelike stimuli used for testing were divided in five sets of 16 pictures. Nonface objects were divided in one set of 350 stimuli always used as base stimuli, and five sets of 16 stimuli for sequences containing only nonface objects (see below). Stimulus resolution and size, screen parameters, and viewing distance were identical to Experiment 1.

#### 2.2.3. Procedure

The procedure was adapted from a recent EEG frequency-tagging study investigating face categorization at various stimulus durations (Retter et al., 2020). Images were presented without inter-stimulus interval (forward and backward-masking) at five stimulation frequencies depending on the sequence: 60 Hz, 30 Hz, 20 Hz, 15 Hz and 12 Hz (i.e., stimulus-onset asynchrony = stimulus duration: 17 ms, 33 ms, 50 ms, 67 ms, 83 ms; Fig. 1C). These frequencies were chosen according to the screen refresh rate (i.e., 60 Hz), such that stimulus durations were 1, 2, 3, 4 or 5 frames. In every sequence, nonface objects were used as base stimuli. Facelike objects were interspersed at 1 Hz in half of the sequences. Thus, for instance, at 60 Hz, facelike objects appeared every 60 images, while at 12 Hz they appeared every 12 images (Fig. 1C). Facelike stimuli were replaced by nonface objects in the other half of the sequences. This led to 10 conditions: 2 categories (facelike objects or nonface objects) × 5 stimulus durations (17 ms, 33 ms, 50 ms, 67 ms, 83 ms; corresponding to 5 stimulation rates: 60 Hz, 30 Hz, 20 Hz, 15Hz, 12 Hz).

Stimulation sequences started with a 0.5-second pre-stimulation interval, followed by a 1.833-second fade-in of increasing contrast modulation depth (0 to 100%). Then, the full-contrast stimulation lasted 15.167 seconds before a 1-second fade-out of decreasing contrast modulation depth (100 to 0%) and a 0.5-second post-stimulation interval. For both facelike and nonface objects interleaved at 1 Hz, each set of 16 images was used in half of the stimulation sequences. The 350 nonface objects used as base stimuli were presented in all sequences. Each experimental condition was repeated 10 times, resulting in 100 sequences throughout the experiment. They were divided in 10 blocks of 10 sequences, each block presenting one sequence per condition. Blocks and sequences within blocks were randomly presented across participants. Stimuli were randomly selected from their respective sets.

#### 2.2.4. Explicit behavioral task

Contrary to Experiment 1, perceptual awareness of illusory faces was expected to vary for each participant, as a function of stimulus duration (i.e., stimulation frequency). Hence, participants were explicitly instructed to attend to the stimuli and to detect facelike objects. After electrode-cap setup, participants were told that rapid sequences of natural images depicting objects will be presented at variable rates and that they will have to report orally after each sequence whether it contained some objects resembling faces. Fifteen facelike images were presented one by one for illustration (not used thereafter). Participants were informed that some sequences will contain several facelike objects and some will not. They were also informed that because images will be presented at rapid rates, false alarms could be frequent. Accordingly, they were asked to report illusory face perception if and only if they noticed several facelike exemplars throughout the sequence. The number of facelike reports after a sequence (out of 10) was submitted to a repeated-measures ANOVA with *Category* (facelike objects *vs*. nonface objects) and *Stimulus duration* (17 ms *vs*. 33 ms *vs*. 50 ms *vs*. 67 ms *vs*. 83 ms) as within-subject factors. Note that this dependent variable corresponds to the number of hits and false alarms for sequences presenting facelike objects and nonface objects, respectively. Mauchly’s test for sphericity violation was performed and Greenhouse-Geisser correction (epsilon: ε) for degrees of freedom was applied whenever sphericity was violated. Post-hoc comparisons were conducted using *T*-tests and the FDR procedure was applied to adjust *p*-values (Benjamini & Hochberg, 1995).

#### 2.2.5. EEG acquisition and preprocessing

EEG acquisition and preprocessing steps were identical to Experiment 1, except for data segmentation into 20-second epochs for each stimulation sequence (100 per participant, 10 conditions × 10 repetitions), including 1 second before the fade-in and 1 second after the fade-out. Following ICA, the mean number of removed components across participants was 2.7 (range: 1–5). The mean number of interpolated channels was 0.5 (range: 0–3).

#### 2.2.6. EEG frequency-domain analysis

As in Experiment 1, the facelike categorization response was tagged at 1 Hz and harmonics. In contrast, contrary to Experiment 1, the general visual response was tagged at different frequencies depending on the sequence (i.e., 60 Hz, 30 Hz, 20 Hz, 15 Hz, 12 Hz and their respective harmonics). For each participant, the 10 epochs recorded for each condition were averaged in the time domain, leading to a single 20-second segment per condition. Epochs were cropped from the onset of the full-contrast stimulation to 16 seconds (4096 time bins). An FFT was applied and amplitude spectra were extracted with a frequency resolution of 1/16 = 0.0625 Hz, leading to 16 frequency bins every 1-Hz step.

For the categorization response, harmonics were included up to the 26^th^ harmonic (i.e., 26 Hz) according to Experiment 1. For the general visual response elicited at variable frequencies up to 60 Hz, we considered harmonics until this frequency for each condition. In other words, only the first harmonic (i.e., 60 Hz) was included for the general response to a 60 Hz-stimulation stream, two harmonics (i.e., 30 Hz and 60 Hz) for the response to a 30 Hz-stimulation, three harmonics (i.e., 20 Hz, 40 Hz, 60 Hz) for a 20 Hz-stimulation, four harmonics (i.e., 15 Hz, 30 Hz, 45 Hz, 60 Hz) for a 15 Hz-stimulation and five harmonics (i.e., 12 Hz, 24 Hz, 36 Hz, 48 Hz, 60 Hz) for a 12 Hz-stimulation. The overall responses were summed across harmonics, excluding those corresponding to the stimulation frequencies and their harmonics (i.e., 12 Hz, 15 Hz, 20 Hz, 24 Hz) for the categorization response. The general and categorization responses will refer to these summed amplitudes thereafter.

The magnitude of each brain response was quantified in mean corrected amplitudes within the ROIs defined in Experiment 1 (i.e., right and left occipito-temporal [r/lOT: P10/9, PO8/7, P8/7] for the categorization response and middle occipital [PO8, O2, Oz, O1] for the general response). To estimate the baseline noise level in a similar frequency range as in Experiment 1 (≈ ± 0.3 Hz), we considered the mean amplitude of 6 surrounding frequency bins (3 on each side), excluding the adjacent bins and the most extreme (minimum and maximum) bins. Statistical analyses were carried on individual corrected amplitudes separately for the categorization and general responses using repeated-measures ANOVAs with *Category* (facelike objects *vs*. nonface objects) and *Stimulus duration* (17 ms *vs*. 33 ms *vs*. 50 ms *vs*. 67 ms *vs*. 83 ms) as within-subject factors. The factor *Hemisphere* (rOT *vs*. lOT) was also included for the categorization response. Mauchly’s test for sphericity violation was performed and Green-house-Geisser correction (epsilon: ε) for degrees of freedom was applied whenever sphericity was violated. Post-hoc comparisons were conducted using *T*-tests with FDR-adjusted *p*-values (Benjamini & Hochberg, 1995). As in Experiment 1, significance of the grand-averaged brain responses was estimated by determining whether the 95% *CI* around the mean amplitude did not include zero. Grand-averaged corrected amplitudes were normalized by their scalp-wide power (McCarthy & Wood, 1985) to illustrate the topography of each brain response regardless of its magnitude.

To evaluate the relationship between the behavioral report of illusory faces and the amplitude of the facelike categorization response, we computed Pearson’s correlations between individual data for both measures and for each stimulus duration. We first divided each measure by its value at the longest duration (i.e., 83 ms) to correct for individual differences in ceiling-level responses (Retter et al., 2020). In other words, this measurement is free from the between-subject variability observed when performance is at ceiling. We then calculated correlations for the four remaining durations. We computed the same correlation for the average of the two shortest durations (i.e., 17 and 33 ms), which lead to mid-level amplitude of the categorization response and mid-level number of facelike reports, and did the same for the two (i.e., 50 and 67 ms) following durations, which lead to larger neural responses and near-ceiling behavioral performance.

In a last step, we determined whether the facelike categorization response emerges as a function of participants’ report of illusory faces for the combination of the two shortest durations (i.e., 17 ms and 33 ms) for which behavioral responses are not at ceiling. For each participant, we averaged preprocessed epochs (i.e., in the time domain) across durations separately for reported (i.e. *aware*) and unreported (i.e., *unaware*) sequences. Following processing steps were identical as in the main analysis to obtain one brain response for perceptually aware sequences, and one response for unaware sequences. Individual summed corrected amplitudes were submitted to a repeated-measures ANOVA with *Awareness* (aware *vs*. unaware) and *Hemisphere* (rOT *vs*. lOT) as within-subject factors. Significance of the grand-averaged brain responses was estimated by determining whether the 95% *CI* did not include zero. For illustration, the difference between aware and unaware sequences was also computed for each participant. Finally, we conducted the same analysis with 12 participants out of 22 to control for a potential influence of the number of sequences associated with awareness or not, which is not equivalent across durations (see Fig. S2).

## 3. Results

### 3.1. Experiment 1: Characterizing the conscious categorization of illusory faces

At 1 Hz and harmonics, we identified two brain responses reflecting the categorization of human faces and facelike objects from variable natural images (Fig. 2A). Summed across harmonics, both categorization responses are of high signal-to-noise ratio (SNR ≈ 3 and 1.5 respectively for human faces and facelike objects; i.e., 200% and 50% of signal increase) and significantly above noise level (i.e., 95% confidence intervals (*CI*) do not include 0; Fig. 2B), despite a larger response to human faces (3.44 ± 0.45 (95% *CI*) μV) than facelike objects (0.72 ± 0.19 μV; 21% of the face categorization response; *F*1,26 = 175, *p* < .001, *η_p_^2^* = 0.87). Importantly, the two categorization responses present close topographies over the occipito-temporal cortex and a right hemisphere advantage (main effect of *Hemisphere: F*1,26 = 7.77, *p* = .009, *η_p_^2^* = 0.23). The human face categorization response is about 23% larger over the right (rOT; 3.78 ± 0.55 μV) than the left occipito-temporal region (lOT; 3.09 ± 0.48 μV) while the facelike categorization response is about 66% larger over rOT (0.91 ± 0.31 μV) than lOT (0.55 ± 0.21 μV). Considering the categorization responses over the whole scalp, about 11.6% and 16.4% of the human face and facelike categorization responses, respectively, are concentrated over rOT (representing less than 5% of the overall channels, i.e., 3 out of 64).

Using channels as observations, we confirmed the similar topographies of the two categorization responses (i.e., highly correlated; *R* = 0.92). In comparison, the correlation between the facelike categorization response and the more central general visual response (*R* = 0.64) is significantly lower (*p* < .001). By splitting EEG data according to stimulus sets (see *Materials and Methods*) and computing correlations between the responses obtained for each set, we also observed that both categorization responses to human faces (*R* = 0.99) and facelike objects (*R* = 0.91) are highly reliable across measurements.

It is worth noting that the visual categorization responses to human and illusory faces are automatically elicited since participants did not explicitly process the two categories but performed an orthogonal cross-detection task with high accuracy (99.3 ± 0.5%) and speed (406 ± 11 ms) without any difference between categories (both *F*s < 1). Similarly, the general visual response recorded at 6 Hz and harmonics over the middle occipital cortex (Fig. 2A & 2C) is not modulated by the stimulus category presented at 1 Hz (human faces: 3.24 ± 0.53 μV; facelike objects: 3.34 ± 0.57 μV; *F* < 1).

Critically, after differentiating participants who overtly reported facelike objects (perceptually *aware* participants, *N* = 13) and those who did not (*unaware* participants, *N* = 14), the categorization response to facelike objects (Fig. 3A) is 153% larger for aware (1.06 ± 0.22 μV) than unaware participants (0.42 ± 0.20 μV; *F*1,25 = 21.3, *p* < .001, *η_P_^2^* = 0.46). In contrast, the human face categorization response is not significantly different between participants (*F* < 1), albeit descriptively larger for aware (3.61 ± 0.73 μV; +10.5%) than unaware participants (3.27 ± 0.63 μV). Likewise, the amplitude of the general visual response (Fig. 3B) does not differ between participants (aware: 3.34 ± 0.85 μV; unaware: 3.25 ± 0.82 μV; *F* < 1), who also do not differ at the behavioral cross-detection task (aware: 99.5 ± 0.4%, 406 ± 15 ms; unaware: 99.1 ± 0.9%, 406 ± 17 ms; both *F*s < 1).

**Figure 3.**
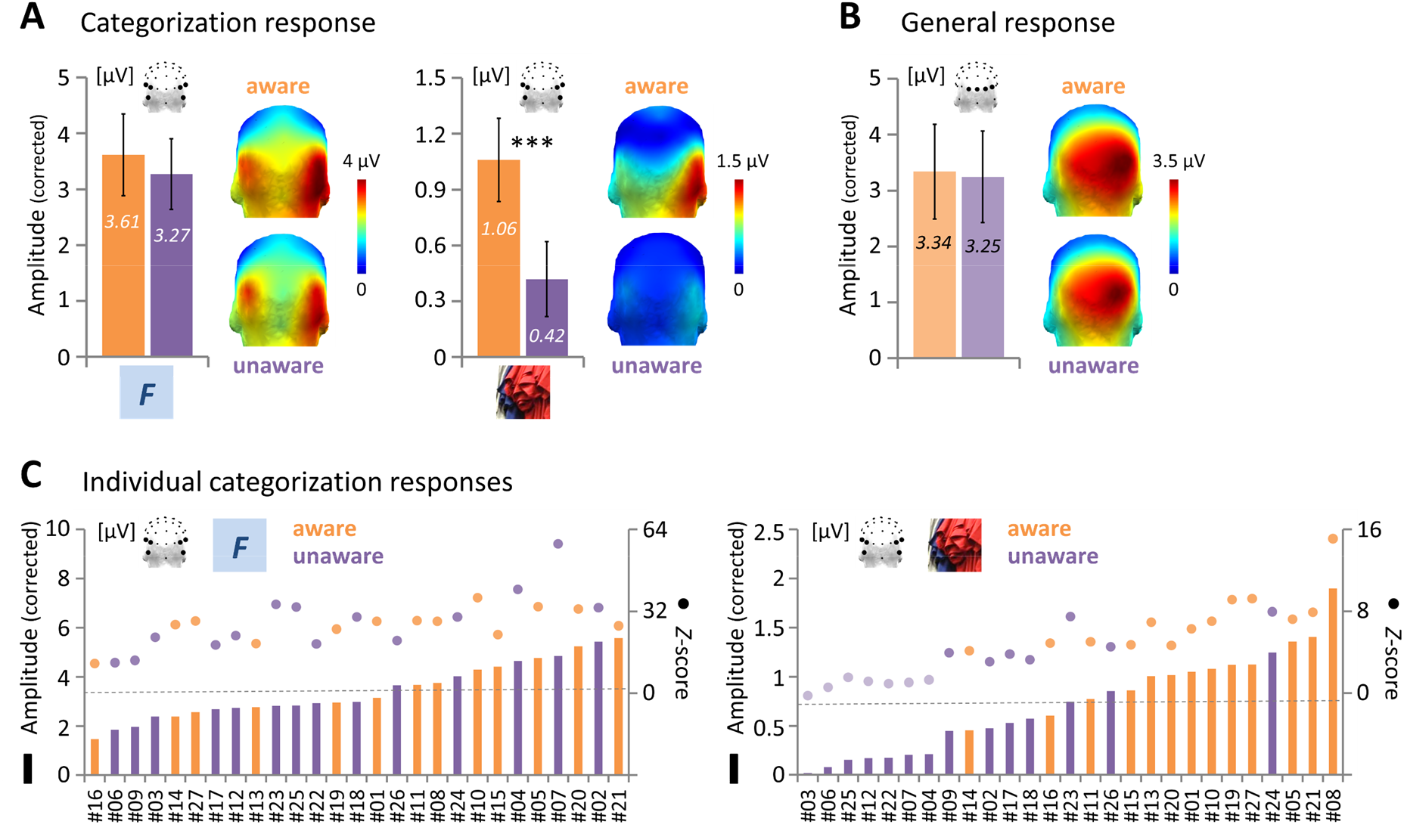
Categorization and general responses according to the perceptual awareness of illusory faces. **A**. Grand-averaged summed corrected amplitude of the categorization responses to human faces (left) and facelike objects (right) averaged across left and right occipito-temporal regions for the perceptually aware (orange) and unaware (purple) participants (*** *p* < .001). **B**. Grand-averaged summed corrected amplitude of the general response over the middle occipital region for the perceptually aware (orange) and unaware (purple) participants (categories collapsed). For both categorization (A) and general (B) responses, topographies are illustrated by head maps (posterior view) and error bars represent 95% confidence intervals. **C**. Individual categorization responses to human (left) and illusory (right) faces for perceptually aware (orange) and unaware (purple) participants. Bars and dots respectively depict summed corrected amplitudes (ranked in ascending order) and *Z*-scores. Lighter dots represent non-significant *Z*-scores (*Z* < 1.64, *p* > .05, one-tailed, signal > noise). The dashed grey line depicts the mean corrected amplitude across all participants.

Next, we estimated the significance of individual categorization responses (Fig. 3C) using *Z*-scores contrasting the amplitude of the response from surrounding noise level (*Z* > 1.64, *p* < .05, one-tailed, signal > noise). Every individual participant presents a strongly significant categorization response to human faces (all *Z*s > 11.64, all *p*s < .001). In contrast, while the categorization response to illusory faces is significant in every perceptually aware participant (all *Z*s > 4.15, all *p*s < .001), it is significant in only half of the unaware participants (i.e., 7 out 14, all *Z*s > 3.07, all *p*s < .002, other half: from *Z* = –0.24, *p* = .59 to *Z* = 1.53, *p* = .063). By ranking the amplitude of individual categorization responses, we observed that 11 out of the 13 largest responses to facelike objects (i.e., above both the mean and the median responses) belong to perceptually aware participants. In other words, with this criterion, EEG data predicts well above chance (*p* = .011) whether a given participant consciously perceives illusory faces (accuracy = 85%). In contrast, predictability is not above chance (*p* = .29) if based on the response to human faces (8 out of the 13 largest responses (62%) belong to perceptually aware participants).

### 3.2. Experiment 2: Perceived vs. unperceived illusory faces in a single brain

Experiment 1 demonstrates that variable objects resembling faces are categorized as faces by the human brain, in association with participants’ report of face pareidolia. However, given that participants were differentiated *a posteriori* from this single report, we conducted Experiment 2 to directly manipulate awareness in a within-subject design and compare the brain responses to perceived and unperceived facelike objects in each participant. After being presented with examples of facelike stimuli, another 22 participants were explicitly instructed to report if they perceived illusory faces after each of a hundred 16-sec-long sequences. Half of the sequences presented facelike objects among nonface objects and the other half presented only nonface objects. Facelike stimuli were always displayed at 1 Hz, but they lasted 17 ms, 33 ms, 50 ms, 67 ms or 83 ms depending on the sequence (i.e., 5 stimulation frequencies: 60 Hz, 30 Hz, 20 Hz, 15 Hz and 12 Hz; Fig. 1C) to make the conscious perception of illusory faces challenging and dissociate sequences associated with perceptual awareness or not.

For sequences containing facelike objects (Fig. 4A), the mean number of facelike reports is low at 17 ms (4.8 ± 1.4 (95% *CI*) reports out of 10) and then increases at 33 ms (7.0 ± 0.9 reports) to reach near-ceiling accuracy from 50 ms (9.0 ± 0.6 reports) to 83 ms (9.6 ± 0.4 reports) with no difference between the three longest durations (all *p*s > .05). In contrast, for sequences containing only nonface objects, facelike reports are very low at all durations (from 2.2 ± 1.2 reports at 17 ms to 1.0 ± 0.5 reports at 83 ms, significant difference between 50 ms and 67 ms, *p* = .044). Hence, the number of accurate facelike reports (i.e., hits) is always greater than the number of erroneous perceptions (i.e., false alarms), even at the shortest 17-ms duration (main effect of *Category: F*1,21 = 238, *p* < .001, *η_p_^2^* = 0.92). In other words, despite the very high constraints put on the visual system at the highest stimulation rate, participants are able to tell apart the two types of sequences (i.e., with or without facelike objects), this difference between the number of hits and false alarms increasing as a function of stimulus duration (*Category* × *Stimulus duration* interaction: *F*2.5,52.3 = 37.5, ε = 0.62, *p* < .001, *η_p_^2^* = 0.64).

**Figure 4.**
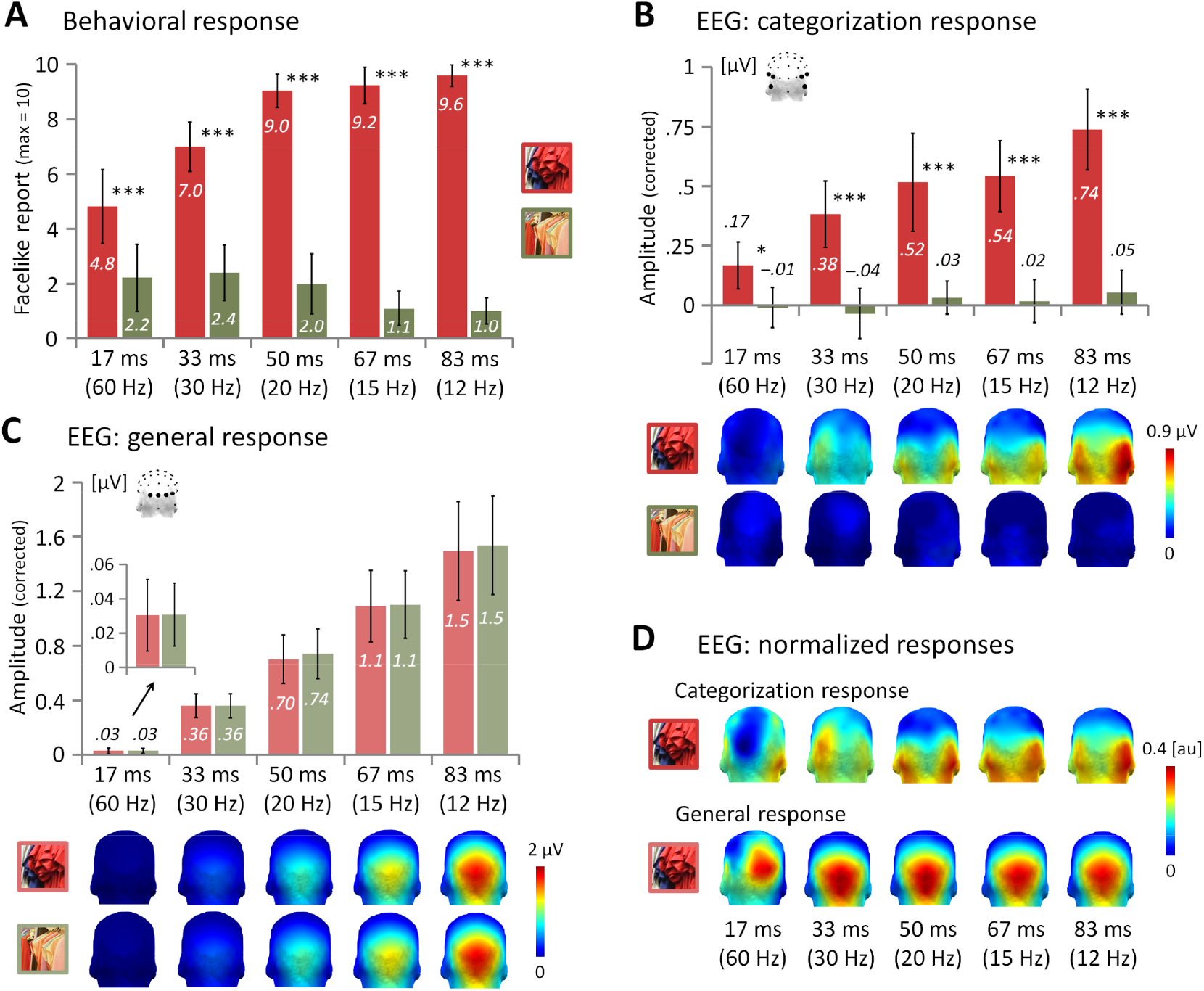
Behavioral and neural responses according to stimulus category and duration. Mean number of facelike reports out of 10 (**A**), grand-averaged summed corrected amplitude of the categorization response over occipito-temporal regions (**B**) and of the general response over the middle occipital region (**C**) for sequences presenting facelike objects (red) or only nonface objects (green) and for the five stimulus durations (* *p* < .05, *** *p* < .001, error bars represent 95% confidence intervals). For both categorization (B) and general (C) responses, topographies are illustrated by head maps (posterior view). **D**. Topographical head maps of normalized categorization and general responses for sequences containing facelike objects illustrate their spatial distribution across stimulus durations (au: arbitrary unit).

At the neural level, the response measured at 1 Hz and harmonics over occipito-temporal regions (Fig. 4B) is also always larger for sequences containing facelike stimuli than only nonface objects (main effect of *Category: F*1,21 = 57.8, *p* < .001, *η_p_^2^* = 0.73). Albeit low at the 17-ms stimulus duration (0.17 ± 0.09 μV), the response to facelike objects is greater than noise level (i.e., 95% *CI* does not include 0) and increases with stimulus duration (until 0.74 ± 0.17 μV at 83 ms), while there is no response to nonface objects (mean amplitude across durations: 0.01 ± 0.05 μV). The difference between the responses to facelike objects and nonface objects thereby increases with duration (*Category* × *Stimulus duration* interaction: *F*4,84 = 6.64, *p* < .001, *η_p_^2^* = 0.24). A right hemisphere advantage is visible at most durations for the categorization response to illusory faces (Fig. 4D), but it does not reach significance (*F* < 1). Like the facelike categorization response, the general visual response (Fig. 4C) is larger than noise at all durations (lowest amplitude: 0.03 ± 0.02 μV at 17 ms) and increases with stimulus duration (until 1.5 ± 0.36 μV at 83 ms, *F*1.4,29.7 = 57.5, ε = 0.35, *p* < .001, *η_p_^2^* = 0.73). However, this middle occipital activity (Fig. 4D) elicited by the rapid stream of stimuli is not different between sequences containing facelike and nonface objects (*F*1,21 = 1.59, *p* = .22).

The increase of both neural and behavioral responses to facelike objects as stimulus duration increases suggests a relationship between these measures, in line with participants’ awareness of face pareidolia. Thus, we evaluated whether participants presenting with larger facelike categorization responses report facelike stimuli in more sequences after weighting both responses by their value at the longest 83-ms duration to correct for ceiling-level neural activity and behavioral performance. For each duration individually, marginal relationships are observed only at 17 ms (*R* = 0.39, *p* = .078) and 33 ms (*R* = 0.42, *p* = .053; other durations: all *R*s < 0.22, all *p*s > .32). When these two shortest durations are combined, neural and behavioral responses become significantly associated (*R* = 0.53, *p* = .011; Fig. 5A). In contrast, there is no association between the two measures for the combination of the two following durations (i.e., 50 and 67 ms; *R* = –0.01, *p* = .97), likely due to the near-ceiling accuracy observed for a majority of participants at these longer durations. In sum, for the two most challenging stimulus durations, which lead to more variable behavioral performance across participants, the overt report of illusory face perception is related to the amplitude of the facelike categorization response.

**Figure 5.**
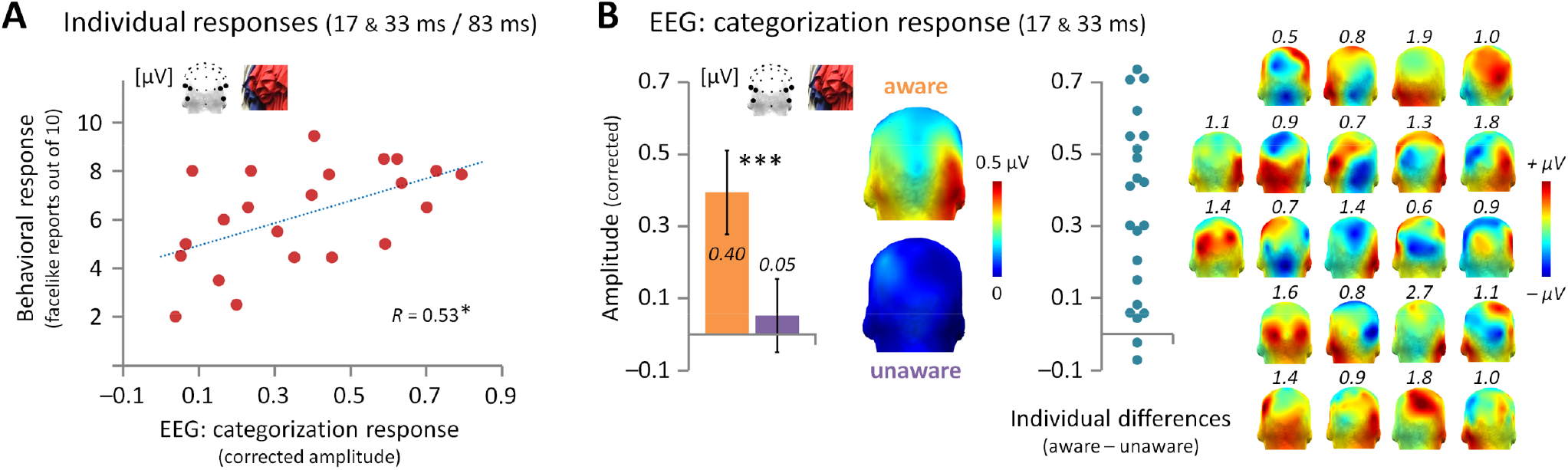
The facelike categorization response predicts the conscious perception of an illusory face. **A**. Correlation between individual summed corrected amplitudes of the facelike categorization response over occipitotemporal regions and the number of facelike reports weighted by their ceiling-level value at the 83-ms duration for the combination of the 17- and 33-ms durations (* *p* < .05). **B**. Left: grand-averaged summed corrected amplitude of the facelike categorization response over occipito-temporal regions depending on participants’ report of illusory faces (perceptually aware sequences: orange, unaware sequences: purple; *** *p* < .001) for the combination of the 17- and 33-ms durations. Error bars represent 95% confidence intervals. Topographies are illustrated by head maps (posterior view). Right: dots depict individual differences between the facelike categorization responses to reported (aware sequences) and unreported (unaware sequences) face pareidolia. Topographies are illustrated by head maps (posterior view) with each individual scale above the map.

Accordingly, we finally explored whether the advent of the facelike categorization response at the 17- and 33-ms durations directly predicts participants’ report of illusory faces. For each participant, we differentiated the facelike categorization responses between sequences wherein they perceived facelike objects (*aware* sequences) and those wherein they did not (*unaware* sequences). Strikingly, the categorization response to facelike stimuli is significantly above noise level only for perceptually aware sequences (0.40 ± 0.12 μV; Fig. 5B) and leads to a larger neural activity compared to unaware sequences (0.04 ± 0.10 μV; *F*1,21 = 40.2, *p* < .001, *ηp*^2^ = 0.66). Descriptively, 20 out of the 22 participants have a larger categorization response when they report facelike objects, such that the sign of the difference between the two conditions predicts above chance whether a given participant was aware of the illusory faces with an accuracy of 91% (*p* < .001). The topography of this difference reveals an advantage for aware sequences over occipito-temporal scalp regions in every individual participant (Fig. 5B). These observations remain unchanged when only 12 participants are considered to equate the number of sequences associated with perceptual awareness or not across the two durations (Fig. S2). This demonstrates that the conscious perception of an illusory face emerges from neural facelike categorization.

## 4. Discussion

Through two experiments, we identify a brain response to a variety of naturalistic facelike objects contrasted to many nonface objects of the same categories in a fast train of forward- and backward-masked stimuli, which, like the categorization response to genuine human faces, is mainly recorded over right occipito-temporal regions. Importantly, this valid neural measure of rapid and automatic facelike categorization under high visual constraints is isolated in individual participants and predicts perceptual awareness with high accuracy, either between groups of perceptually aware *vs*. unaware participants (Experiment 1), or between stimulation sequences according to participants’ report of face pareidolia (Experiment 2). Hence, thanks to the advantages of EEG frequency-tagging to objectively measure automatic visual categorization in the brain, and capitalizing on a visual illusion to investigate the richness of face perception beyond the mere categorization of human faces, the present study provides a clear signature of face pareidolia in the human brain.

Face pareidolia is one of the most remarkable examples of ubiquitous illusory percept in the human species. It has been widely used by painters (e.g. Giuseppe Arcimboldo, 1527–1593) or photographers (e.g., Robert & Robert, 1996), and more than 70% of pictures represent a face when searching “pareidolia” on the web (estimation made in January 2021 with the first 100 different pictures in Google Images). Accordingly, going beyond previous efforts with scalp EEG and neuroimaging (e.g., Caharel et al., 2013; Churches et al., 2009; Dolan et al., 1997; George et al., 2005; Rossion et al., 2011), the first major achievement of the present study is to provide a rich and valid measure of face pareidolia under the variable viewing conditions in which it takes place in the natural visual environment. The use of various natural views of facelike objects contrasted to equivalent nonface stimuli also makes unlikely the contribution of low physical variability between facelike objects, contrary to stimuli segmented from their background which artificially increases facelikeness by delineating a global face shape (see Davidenko et al., 2012 for a discussion). Naturalistic stimuli also implies figure-ground segregation, an integral part of object perception (Wagemans et al., 2012). In addition, the facelike categorization response reflects spontaneous face pareidolia, that is, the automatic perception (i.e., unintentional and hard to suppress) of an illusory face, at a glance. It is also worth noting that the facelike categorization response is a direct differential response (i.e., without post-hoc subtraction), such that it would have been absent if facelike objects were not discriminated from nonface objects, as observed in Experiment 2 for sequences wherein only nonface objects are displayed. The response is objective (i.e., recorded at a pre-experimentally defined frequency and its harmonics), and highly sensitive and reliable, as estimated in Experiment 1. These properties are critical to identify unambiguous individual brain responses, estimate their significance, and relate them to participants’ report of face pareidolia.

In Experiment 1, we also clarify how similar is the categorization of illusory faces to the categorization of genuine human faces, both quantitatively and qualitatively. Consistent with previous studies using the same paradigm, we observe that the categorization of human faces elicits a large response of about 4 μV over the occipito-temporal cortex, with a right hemisphere advantage (e.g., Jacques et al., 2016; Retter & Rossion, 2016). The facelike categorization response has a similar topography with an increased right-hemispheric dominance, but is of about 20% of the face categorization response in amplitude overall (35% when considering only perceptually aware participants). At least three non-mutually exclusive interpretations can explain this observation. First, the two responses may be generated by the same face-selective regions distributed along the VOTC (Gao et al., 2018; Jonas et al., 2016; Sergent et al., 1992; Zhen et al., 2015; Grill-Spector et al., 2017 for review), with a weaker activation overall for facelike objects. Such diminished responsiveness could be due to the absence of some cues pertaining to human faces in facelike stimuli. For instance, while both shape and color information are important cues for visual recognition (Gegenfurtner & Rieger, 2000; Tanaka et al., 2001), and while color contributes to about 20% of the face categorization response for photographs (Or et al., 2019), color does not inform about facelikeness in facelike objects. A second interpretation may be that only a subset of face-selective regions generates the facelike categorization response. Neuroimaging studies have associated the perception of an illusory face with the lateral part of the middle fusiform gyrus (Andrews & Schluppeck, 2004; Dolan et al., 1997; Kanwisher et al., 1998; McKeeff & Tong, 2007; Rossion et al., 2011), sometimes considered as a key region for the perception of a global face configuration (Andrews et al., 2010; Goffaux et al., 2013; Rossion et al., 2011). Relatedly, the facelike categorization response is strongly right-lateralized, the right hemisphere being particularly involved in the perception of a global facelike configuration (Caharel et al., 2013; Parkin & Williamson, 1987; Rossion et al., 2011). However, despite a numerical advantage, there is no significant hemispheric asymmetry in Experiment 2, which may be due to the explicit instruction to attend to illusory faces increasing the left hemisphere contribution (see Quek et al., 2018). A last interpretation may be that facelike stimuli either elicit a full response within the whole face-selective network when they are perceived as faces, or they do not elicit a face-selective response at all when they are not perceived as faces, leading to a lower response in average. In other words, neural categorization could be strictly identical for facelike objects and human faces, but artificially appear weaker for facelike objects due to more occasional occurrences. Interestingly for our purpose, this account concurs with the view that face categorization emerges all at once from the linear accumulation of sensory evidence and reflects perceptual awareness (Harris et al., 2011; Navajas et al., 2013; Retter et al., 2020; Tong et al., 1998).

In that respect, the second major achievement of our study is to characterize to what extent the neural categorization of facelike objects reflects conscious illusory face perception, extending prior work on the association between a neural response to simple facelike stimuli and their perceptual interpretation as a face (Andrews & Schluppeck, 2004; Bentin et al., 2002; George et al., 2005; Shafto & Pitts, 2015). In Experiment 1, we observe a strong categorization response to facelike objects in participants who report face pareidolia post-stimulation compared to a weak response in those who do not, and reveal that individual facelike categorization responses predict this association. More strikingly, this relationship is confirmed in a single group of participants in Experiment 2, with a significant correlation between the facelike categorization response and the number of illusory face reports. As a result, when directly comparing sequences associated with awareness or not in this experiment, the facelike categorization response is observed only when face pareidolia are reported, with a conspicuous difference between the two types of sequences in each individual participant. Hence, given that the response varies greatly as a function of perceptual awareness despite identical stimuli, and given that these stimuli are directly contrasted to other stimuli depicting the same object categories, our study provides original evidence supporting the view that neural face categorization is a signature of conscious face perception (Harris et al., 2011; Navajas et al., 2013; Retter et al., 2020; Tong et al., 1998).

The strict absence of facelike categorization response to unreported face pareidolia in Experiment 2 points toward an all-or-none neural categorization function in response to sensory evidence gradually accumulating in early visual areas, as mentioned above. However, in Experiment 1, although the large difference between perceptually aware and unaware participants indicates that the bulk of the response reflects conscious illusory face perception, the response is not completely abolished in unaware participants. This suggests that a residual selective response to facelike objects could be observed in the absence of overt report. One explanation may be that some cues elicit a differential response between facelike and nonface objects, albeit non-sufficient to trigger full categorization. For instance, facelike objects all depict “eyelike” or “mouthlike” features (e.g., Fig. 1A), sometimes considered critical features to perceive a nonface stimulus as a face (e.g., Omer et al., 2019). Similarly, some image statistics covary with facelikeness, such as more contrast (i.e., higher spatial frequencies) in the upper part of the image. This visual property is well-known to already attract attention in newborns, as a precursor to develop face perception (e.g., Johnson et al., 1991; Simion & Di Giorgio, 2015 for review). Moreover, the presence of human faces in this experiment could have primed face-related cues in facelike stimuli. Alternatively, since perceptually aware and unaware participants in Experiment 1 were differentiated *a posteriori* following a single awareness assessment that summarizes the experience of almost 3000 stimulation cycles including more than 250 facelike stimuli, the small response in unaware participants could be due to an idiosyncratic confound such as the criteria to define an illusory face or the ability to remember sparse occurrences of face pareidolia. Therefore, the number of participants who consciously perceived illusory faces may have been underestimated in this experiment. In this context, it should be noted that both the general visual response elicited by the rapid stream of stimulation and the efficiency at the cross-detection task do not differ between participants, making unlikely the contribution of visual attention. Irrespective of these potential limitations, they do not concern Experiment 2, which reveals a striking difference between perceptually aware and unaware responses to facelike objects in a single group of participants.

By manipulating stimulus duration while participants explicitly process facelike stimuli, Experiment 2 additionally provides important information about the optimal conditions for face pareidolia to arise within a fast train of forward- and backward-masked stimuli. Albeit low at the shortest 17-ms duration, both facelike reports and the categorization response are higher for sequences containing facelike objects than only nonface objects at every duration. This means that participants are already able to perceive some illusory faces at 17 ms, in agreement with a minimal duration of approximately 13-17 ms to behaviorally or neurally categorize human faces in rapid streams of natural images (Keysers et al., 2001; Retter et al., 2020), or other visual objects in various experimental designs (Bacon-Macé et al., 2005; Fisch et al., 2009; Mohsenzadeh et al., 2018; Potter et al., 2014). Facelike reports then increase at 33 ms and reach ceiling at 50 ms. In contrast, the amplitude of the facelike categorization response increases until the longest 83-ms duration and could still increase beyond (e.g., a larger response is observed with a 167-ms duration for perceptually aware participants in Experiment 1), unlike the response to human faces, which saturates at 83 ms (Retter et al., 2020). Moreover, behavioral and neural responses become decorrelated at 50 ms and 67 ms. This dissociation between behavioral and neural responses from 50 ms is not surprising given that one behavioral response was recorded after each sequence presenting at least 192 stimuli and sometimes including 16 facelike objects. Therefore, participants could have reported illusory face perception from a few facelike stimuli, such that accuracy rapidly reached ceiling at the intermediate 50-ms duration, but the number of categorized stimuli within a sequence could still increase beyond 50 ms. This accords with previous studies showing that various factors, such as the overlap of sensory information with forward and backward stimuli, lead to the categorization of only a fraction of stimuli at short presentation times (e.g., Bacon-Macé et al., 2005). Interestingly, the number of false alarms only decreases from 67 ms, and is significantly higher than zero at all durations. Thus, the explicit instruction to detect facelike stimuli among other stimuli within rapid sequences depicting many object categories makes participants incline to falsely report facelike objects, even at ceiling-level stimulus durations. Importantly, however, false alarms are not associated with a significant neural categorization response. Since erroneous perceptions can be driven by any stimulus within a sequence of nonface stimuli, they do not occur periodically and do not translate at 1 Hz and harmonics in the EEG spectrum.

In sum, by using a widely variable set of naturalistic facelike objects contrasted to another variable set of the same object categories, we measure a rich neural categorization response to the facelike stimuli that is intimately related to their conscious perception as faces. Hence, coupling face pareidolia, which reflects the high inclination of the human visual system to perceive faces beyond genuine faces, and EEG frequency-tagging, which measures rapid categorization in the brain with objectivity, sensitivity, reliability and validity, provides a powerful approach to characterize the neural underpinnings of conscious face perception. In doing so, we corroborate the view that the perceptual awareness of a face emerges from a categorical response to unconscious sensory inputs (Harris et al., 2011; Navajas et al., 2013; Retter et al., 2020; Tong et al., 1998). More generally, since visual categorization is subtended by a set of category-selective regions in the VOTC, as shown with the present paradigm in fMRI (Gao et al., 2018) and human intracerebral recordings (Hagen et al., 2020; Jonas et al., 2016), it is tempting to consider these regions as neural substrates of perceptual awareness, a long-standing issue in cognitive neuroscience (Boly et al., 2017; Dehaene et al., 2017; Koch et al., 2016; Odegaard et al., 2017 for reviews). We acknowledge that much research must be carried out to further clarify this issue.

## Acknowledgments

The authors are grateful to Romain Patrux and Lucas Ronat for their help in data collection. This work was supported by grants from the “Conseil Régional Bourgogne Franche-Comté” (PARI grant), the FEDER (European Funding for Regional Economic Development), the French “Investissements d’Avenir” program (project ISITE-BFC, contract ANR-15-IDEX-0003), and the French National Research Agency (ANR, contract ANR-19-CE28-0009).

