## Supplement for "Rapid neural categorization of facelike objects predicts the perceptual awareness of a face (*face pareidolia*)"

### *Supplementary information*

### Supplementary Materials and Methods

#### *Stimulus selection*

We used a variety of biological[1] and manufactured objects[2] as nonface stimuli, including between 3 and 20 exemplars for each category. Facelike stimuli matched several object categories[3], with 1 to 5 exemplars for each category. Facelike stimuli were selected among 224 images collected from the Internet when searching for 'face pareidolia'. Selection was made according to the images judged as the most facelike in a pretest also including 224 nonface images selected from the main set. We told 142 participants (111 females, 15 left-handed (12 females), mean age: 18.9 ± 1.5 years, range: 17–26 years) that natural images depicting objects will be presented with some of them resembling faces. In a two forced-choice facelike categorization task (i.e., facelike *vs.* non-facelike), participants responded as fast as possible for each image (i.e., 464 trials) using two keys with their left and right index fingers (dominant finger for the facelike response). Each trial began with a fixation cross displayed at the center of a monitor for 467 ms, followed by the test image for 133 ms (Fig. S1). Hence, while stimuli were not forward- and backward-masked as in the main experiment, they were presented for a brief duration that constrained participants to judge facelikeness at first glance. Stimuli were followed by an inter-trial interval of 1.2 s. Facelike and nonface images were presented randomly in 8 blocks of 56 trials. For each facelike image, we calculated the percentage of facelike responses across participants and mean response time (RT, only for facelike responses ranging between 100 and 1000 ms). In average across facelike images, the percentage of facelike responses was 83.5 ± 14.3% (*SD*) (range: 23% – 96%) and mean RT was 473 ± 33 ms (range: 416 – 621 ms). To combine both measures and exclude speed-accuracy trade-offs, we calculated inverse efficiency (i.e., RT divided by accuracy) for each image. It ranged from 445 (i.e., judged as the most facelike) to 2299 (i.e., judged as the less facelike). Based on these data, we selected the 86 facelike images judged as the most facelike (inverse efficiency range: 445 – 508).

---

[1] birds, cats, cells, dogs, eggs, flowers, fruits, horses, plants, trees, vegetables.

[2] bags, bells, belts, blocks, bowls, boxes, brushes, cameras, candies, canoes, car parts, casings, chairs, clocks, clothes, cookers, crates, cups, electric devices, glasses, graters, guitars, houses, jars, lamps, latches, lids, mail boxes, metallic devices, pant pockets, pastries, pipes, plaques, plastic devices, plates, robots, scooters, spoons, staplers, taps, telephones, trashes, washing machines, toilets, yoghurts.

[3] bags, bells, belts, blocks, bowls, boxes, brushes, candies, canoes, car parts, casings, cells, clocks, clothes, cookers, crates, cups, electric devices, eggs, fruits, glasses, graters, houses, jars, latches, lids, mail boxes, metallic devices, pant pockets, pastries, pipes, plants, plaques, plastic devices, plates, robots, scooters, spoons, staplers, taps, trashes, trees, vegetables, washing machines, toilets, yoghurts.

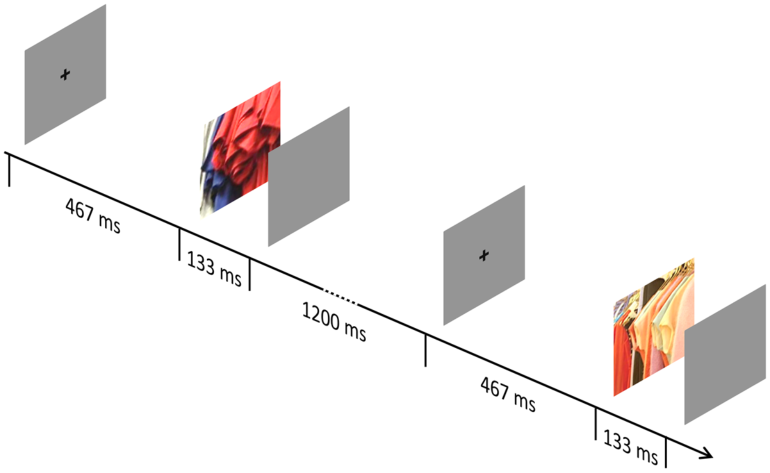


**Figure S1. Experimental design for facelike stimulus selection.** Participants were presented with 464 trials starting with a fixation cross for 467 ms, followed by the stimulus for 133 ms and an inter-trial interval of 1200 ms. Participants were instructed to categorize the stimulus as facelike or non-facelike as fast as possible after stimulus-onset.

**Table S1. Significance of the frequency-tagged responses and their harmonics (i.e., integer multiples).** To define the general (6 Hz and harmonics) and categorization (1 Hz and harmonics) responses in Experiment 1, we determined the range of significant harmonics using *Z*-scores (amplitude at the target frequency minus mean amplitude of the surrounding noise (20 frequency bins, 10 on each side, excluding the immediately adjacent and the 2 most extreme bins) divided by the standard deviation of the noise) calculated on the mean EEG amplitude spectrum across channels and participants for sequences containing human faces. Significant Z-scores ($Z > 1.64$, $p < .05$, one-tailed, signal > noise) are indicated in bold. Harmonics were considered until Z-scores were no longer significant for both the general (blue) and categorization (red) responses.

| Frequency [Hz] | *Z*-score | Frequency [Hz] | *Z*-score |
|---|---|---|---|
| 1 | **16.4** | 25 | **2.33** |
| 2 | **49.9** | 26 | **4.33** |
| 3 | **53.5** | 27 | 1.56 |
| 4 | **24.8** | 28 | -0.23 |
| 5 | **21.8** | 29 | **4.31** |
| 6 | **198** | 30 | **34.9** |
| 7 | **37.8** | 31 | -0.76 |
| 8 | **33.4** | 32 | 1.57 |
| 9 | **30.8** | 33 | **2.06** |
| 10 | **18.5** | 34 | 1.33 |
| 11 | **15.4** | 35 | 0.13 |
| 12 | **58.1** | 36 | **22.9** |
| 13 | **13.9** | 37 | -0.29 |
| 14 | **9.33** | 38 | 0.04 |
| 15 | **9.12** | 39 | -1.37 |
| 16 | **9.23** | 40 | **1.67** |
| 17 | **6.92** | 41 | **2.32** |
| 18 | **80.8** | 42 | **25.7** |
| 19 | **7.29** | 43 | -0.28 |
| 20 | **3.04** | 44 | **1.75** |
| 21 | **2.52** | 45 | 0.29 |
| 22 | **3.16** | 46 | **1.84** |
| 23 | **2.87** | 47 | 0.64 |
| 24 | **31.5** | 48 | **16.7** |

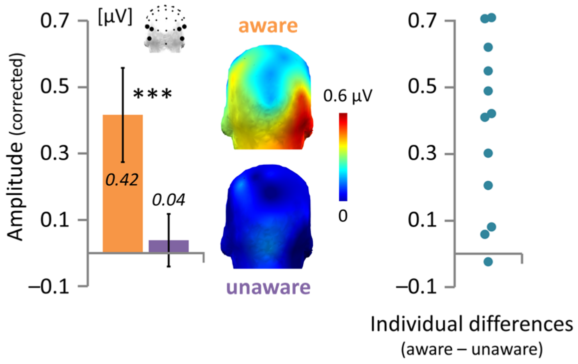


**Figure S2. Controlling for the number of sequences in the relationship between neural facelike categorization and face pareidolia.** Analysis of the whole sample of participants reveals that the facelike categorization response for the combination of the two shortest durations (i.e., 17 ms and 33 ms) emerges only when participants report face pareidolia (i.e., for perceptually aware sequences; see *Results* and Fig. 5B). However, since the total number of perceptually aware sequences is greater for the 33-ms than the 17-ms duration (154 *vs.* 106), leading to a greater ratio between aware and unaware sequences for the 33-ms than the 17-ms duration (0.93 *vs.* 2.33; see also *Results* and Fig. 4A for the mean number of aware sequences (i.e., facelike reports) at each duration), we conducted the same analysis by considering only 12 participants out of 22 to equate the number of perceptually aware sequences across durations (= 69), and to reduce the difference between the number of aware and unaware sequences for each duration (i.e., 69 *vs.* 51 for each duration; $\chi^2 = 2.7$, $p = .10$). This analysis confirms that the facelike categorization response over occipito-temporal regions is related to participants' report of illusory faces, with a significant response only for aware sequences (0.42 ± 0.14 (95% *CI*) µV, orange), which is larger than the response for unaware sequences (0.04 ± 0.08 µV, purple; *** $F_{1,11} = 26.5$, $p < .001$, $\eta_p^2 = 0.71$; error bars represent 95% confidence intervals; topographies are illustrated by head maps with a posterior view). Individual differences between the facelike categorization responses for perceptually aware and unaware sequences show that 11 out of the 12 participants have a larger response for aware sequences (perceptual awareness is predicted by the sign of the difference; accuracy: 92%, $p < .004$).
